# Translational neurophysiological biomarkers of N-methyl-D-aspartate receptor dysfunction in serine racemase knockout mice

**DOI:** 10.1101/2020.04.01.020438

**Authors:** Andrea Balla, Stephen Ginsberg, Atheir I. Abbas, Henry Sershen, Daniel C. Javitt

## Abstract

Alterations in glutamatergic function are well established in schizophrenia (Sz), but new treatment development is hampered by the lack of translational pathophysiological and target engagement biomarkers as well as by the lack of animal models that recapitulate the pathophysiological features of Sz. Here, we evaluated the rodent auditory steady state response (ASSR) and long-latency auditory event-related potential (aERP) as potential translational markers. These biomarkers were assessed for their sensitivity to both the N-methyl-D-aspartate receptor (NMDAR) antagonist phencyclidine (PCP) and to knock-out (KO) of Serine Racemase (SR), which is known to lead to Sz-like alterations in function of parvalbumin (PV)-type cortical interneurons. Both PCP and SRKO led to significant increases of ASSR, consistent with PV interneuron effects. Similar effects were observed in mice with selective NMDAR KO on PV interneurons. By contrast, PCP but not SRKO reduced the amplitude of the rodent analog of the human N1 potential. Overall, these findings support use of rodent ASSR and long-latency aERP, along with previously described measures such as mismatch negativity (MMN), as translational biomarkers, and support SRKO mice as a potential rodent model for PV interneuron dysfunction in Sz.

## Introduction

*N*-methyl-D-aspartate-type glutamate receptor (NMDAR) antagonists, such as phencyclidine (PCP) and ketamine, induce a wide range of symptoms and cortical neurocognitive deficits similar to those observed in schizophrenia (Sz), leading to theories of impaired NMDAR function (Coyle, 1996; Javitt and Zukin, 1990, 1991; Krystal et al., 1994). NMDAR are modulated by the endogenous amino acids glycine and D-serine, which bind to an allosteric modulatory site that regulates channel opening (Javitt et al., 1999; Javitt and Zukin, 1989). Moreover, disturbances in both glycine and D-serine metabolism are reported in Sz (rev. in (Kantrowitz and Javitt, 2012)). To date, however, no FDA-approved glutamate-based treatments are available, in part due to lack of translational biomarkers that permit integration across rodent and human models.

In Sz, deficits are observed in a range of neurophysiological biomarkers including auditory mismatch negativity (MMN), steady-state response (ASSR) and N1 refractoriness (rev. in (Javitt et al., submitted; Javitt et al., 2008). We have recently shown that rodent MMN is highly sensitive to both NMDAR antagonists and agonists and thus may potentially be used as both a translational and target engagement biomarker (Lee et al., 2018). Auditory N1 refractoriness and ASSR, however, have been studied to a lesser degree. Here, we evaluated the sensitivity of these measures both to NMDAR antagonist administration and to genetic manipulations involving D-serine synthesis and parvalbumin interneuron function that potentially recapitulate pathophysiological features of Sz, with the goal of evaluating their utility for translational drug development.

In the ASSR paradigm, a series of repetitive clicks are played at rates of typically between 20 and 100 Hz and elicit an entrained response at the stimulation frequency. Human ASSR shows a natural resonance at 40 Hz, consistent with an interaction between glutamatergic principal neurons and local circuit parvalbumin (PV)-type GABAergic interneurons (rev. in (Dienel and Lewis, 2018; Javitt et al., 2008)). Deficits in ASSR were first reported ∼20 yrs ago (Kwon et al., 1999), and have been replicated consistently since that time when relatively short (∼1 sec) interstimulus intervals are used (rev. in (O’Donnell et al., 2013; Tada et al., 2019)). ASSR deficits have been postulated to interrelate with oxidative-stress driven impairments in PV interneuron function (Gonzalez-Burgos et al., 2010; Steullet et al., 2017), although increases have been reported in paradigms that use longer (e.g. ∼3s) intervals (Hamm et al., 2012; Kim et al., 2019).

Deficits are also observed in other aspects of the response, such as phase delay (Kwon et al., 1999; Roach et al., 2019). As opposed to other auditory ERP, such as MMN, ASSR appears to be intact in individuals at clinical high risk for Sz (Lepock et al., 2019; Tada et al., 2016), although it is impaired in chronic stages (rev. in (Tada et al., 2019)). Thus, it may index downregulation of cortical circuits over time during early stages of the disorder and could possibly be a target for early intervention.

In healthy human volunteers, NMDAR antagonist administration is reported to increase gamma band (40-85 Hz) neural response to auditory (Hong et al., 2010; Plourde et al., 1997) or visual (Grent-’t-Jong et al., 2018) stimuli, but to decrease response in delta and theta frequency ranges (Hong et al., 2010). In these studies, increases in gamma were associated with clinical symptoms, supporting their relevance to Sz (Hong et al., 2010). More recently, however, a reduction in ASSR following ketamine administration was reported (Curic et al., 2019). In rodents, both increases (Leishman et al., 2015; Sullivan et al., 2015; Vohs et al., 2012) and decreases (Schuelert et al., 2018; Sivarao et al., 2013) in ASSR are also reported, potentially explainable based on varying degrees of NMDAR occupancy across studies (Sivarao et al., 2016).

In the auditory refractoriness paradigm, individual stimuli (e.g. tones, clicks) are presented repeatedly at intervals of several hundred milliseconds to seconds, with increasing amplitude as a function of interval length. Under these stimulation conditions, the tones elicit a series of long-latency auditory ERP (aERP) components, including the P1, N1 and P2 potentials. Like ASSR, deficits in auditory N1/P2 response have been extensively documented in Sz (Ford et al., 2001; Javitt et al., 2008; Shelley et al., 1999; Turetsky et al., 2009). Also like ASSR, Sz-like deficits in auditory N1 generation are induced by NMDAR antagonists in both monkey (Javitt et al., 2000) and rodent (Connolly et al., 2004; Lee et al., 2018; Maxwell et al., 2006; Schuelert et al., 2018) models. However, as opposed to ASSR, power associated with the auditory N1/P2 component maps predominantly to the theta frequency range, implicating somatostatin (SOM) interneuron related circuits (Javitt et al., 2018; Javitt and Sweet, 2015; Womelsdorf et al., 2014). These measures thus may provide complementary insights into integrity of specific interneuron mechanisms within cortex.

In the present study, we evaluated the integrity of these measures in a putative rodent model of Sz associated with knockout of the serine racemase (SR) gene. SR is the primary enzyme mediating D-serine synthesis in brain. The presence of D-serine in rodent brain was first demonstrated by Hashimoto et al. (Hashimoto et al., 1992) and was subsequently shown to reflect interconversion of L- to D-serine (Dunlop and Neidle, 1997) via SR (Coyle and Balu, 2018; Wolosker et al., 1999). Moreover, expression-reducing deficits in the non-coding region of the serine racemase (*SRR*) gene are reported in Sz (Schizophrenia Working Group of the Psychiatric Genomics, 2014), suggesting potential etiological involvement.

Over recent years, SRKO mice have been extensively phenotyped and have been shown to have the predicted reduction in brain D-serine levels, as well as specific homologies to features of Sz (rev. in (Coyle and Balu, 2018)). These include most prominently a significant reduction in PV interneuron number (Steullet et al., 2017) and loss of cortical grey matter in both frontal (DeVito et al., 2011) and sensory (Balu et al., 2012) cortex.

The present study therefore evaluated the degree to which SRKO mice showed abnormalities in auditory aERP, potentially linked to disturbances in PV interneuron function. aERP were obtained both prior to and following administration of the NMDAR antagonist PCP. We hypothesized that SRKO mice would show abnormalities similar to those induced by acute NMDAR antagonist administration. Finally, in order to obtain a non-pharmacological comparison group, we constructed mice in which the NR1 subunit of the NMDAR (GRIN1) was knocked-out specifically in PV interneurons using a Cre/Lox approach. As with SRKO mice, we hypothesized that this would produce a phenotype similar to that observed for acute NMDAR antagonists.

## Materials and Methods

This study was carried out in accordance with the Guide for the Care and Use of Laboratory Animals as adopted by the National Institutes of Health and approved by Nathan Kline Institute Animal Care and Use Committee. Heterozygous mice mutants (+/-) with a deletion of the gene encoding SR which converts L-Ser to D-Ser (Basu et al., 2009) were initially obtained from Coyle laboratory of McLean Hospital, Harvard Medical School. At least F10 hybrids were bred in house resulting in wildtype (WT) (+/+), heterozygous (HET) (+/-) and homozygous (HOM) (-/-) littermates with consequent reduction of D-serine level by 50% and 90% respectively. The genetic testing was performed by Transnetyx Inc. (Cordova, TN). The animals were maintained under a 12 h/12 h dark/light cycle and were allowed food and water ad libitum.

Approximately 3-4 month old mice underwent stereotaxic implantation of tripolar electrode assemblies (Plastics One Inc., Roanoke, VA) under isoflurane anesthesia. Stereotactic coordinates were AP −2.50 from bregma, ML +3.50mm, and depth 1-2.25 mm (15° angle). Three stainless-steel electrodes, mounted in a single pedestal, were aligned along the sagittal axis of the skull (at 1 mm intervals (positive, ground and negative). Positive electrodes (2.25 mm) were placed above the left primary auditory cortex region.

Negative electrodes (1mm) were placed adjacent to the ipsilateral neocortex. Ground electrodes (1 mm) were located between recording and reference electrodes. The electrode pedestal is secured to the skull with cyanoacrylic cement (PlasticsOne, Roanoke, VA). An analgesic (Buprenex 0.3 mg/kg, s.c.) was administered after surgery and every 24 hours thereafter as needed. Following surgery, animals were individually housed and allowed to recover for at least 7 days prior to the recording of evoked potentials. At the completion of all recordings, mice were anesthetized with ketamine hydrochloride and acepromazine maleate 1:1 mixture (1 μl/g i.p.). The brain was fixed with 10% formaldehyde solution and was stored in 30% glucose solution for determination of placement of the electrodes. A total of 100 animals were used for this study (31 WT (+/+), 33 HET (+/-) and 36 HOM (-/-)).

### Neurophysiology studies

After a recovery period, aERP measures were recorded from freely moving mice in a clear polycarbonate bowl with food ad libitum in a sound proof chamber. Mice were allowed to habituate for an hour to the new surroundings. An initial testing session was performed for each animal to permit assessment of genetic status. Each animal then participated in two additional drug-treatment sessions in random order with at least a 2-week intervening period. Each session consisted of a 2-hour baseline recording, followed by acute administration of either PCP (10 mg/kg) or saline i.p. followed by an additional 2-hour recording.

Stimuli were administered free-field using a speaker located above the recording chamber generated by a Presentation program, calibrated to 80 dB. To assess the N1 refractoriness function mice were exposed to separate blocks of 200 stimuli (6000 KHz 60msec) at 0.5, 1, 3 and 6 s interstimulus interval (ISI). To assess ASSR, 500-msec trains of 0.1 msec clicks were delivered at stimulation rates of 20, 25, 40, 50, and 80 Hz in separate blocks (300 stimuli each) with 500-ms interval between click trains (overall stimulus onset asynchrony = 1 s).

Neuroelectric signals were impedance matched with unity gain pre-amplifiers located near the electrode, and further differentially amplified with an appropriate bandpass (typically 0.3 Hz to 3 kHz). For additional noise reduction we used 50/60 Hz noise eliminator (Hum Bug, Quest Scientific, North Vancouver, CA). Data were acquired continuously along with digital stimulus identification tags at a digitization rate exceeding 10 kHz using a Neuroscan SCAN system. Epoching, sorting, artifact rejection and averaging were then conducted off-line. For ASSR a FFT spectral analysis was performed, highest amplitudes were recorded in the relevant frequency bands.

### PV-specific GRIN1 KO mice

Grin1-/PV mice were generated by crossing female mice with floxed NMDAR (Grin1tm2Stl/J) and male mice with Cre recombinase expressed specifically within PV-interneurons (129P2-Pvalbtm1(cre)Arbr/J mice). Both mouse lines were obtained from Jackson laboratory (Bar Harbor, ME) and were tested relative to mice with the same C57BL/6J background. Genotype was confirmed by tail-clip (Trasnetyx, Inc. Cordova, TN).

### Statistical analysis

Data were analyzed across animals using mixed-model Analysis of Variance (ANOVA). Separate analyses were performed for 1) the initial (non-drug) session from each animal, and 2) pre/post treatment effects from saline/PCP drug-challenge days. Least significant difference (LSD) analyses were used for post-hoc testing All statistics are two-tailed with pre-specified α level of significant set at p<.05.

## Results

For SRKO mice, ASSR and aERP responses were obtained both at pre-treatment and following challenge with PCP 10 mg/kg. For PV-selective GRIN1 KO animals, only ASSR was obtained.

### Auditory Steady State Response

Analyses were performed on both the baseline (500-ms pre-stimulus) and stimulus (0-500 ms) period (**Fig. 1A,B**). During active stimulations, 500-ms click trains were presented at stimulation frequencies of 20, 25, 40, 50, and 80 Hz and peak power was calculated at each frequency. We performed a 2-way mixed-model ANOVA on the initial (non-drug) testing day for each animal to examine the main effects of stimulation frequency and genotype on ASSR peak power as well as any interactions; followed by a 4-way ANOVA on combined data from the two drug challenge days to examine the main effects of stimulation frequency, genotype, drug treatment (saline vs PCP), and pre-or-post-treatment vs ASSR peak power as well as any interactions.

**Figure 1:**
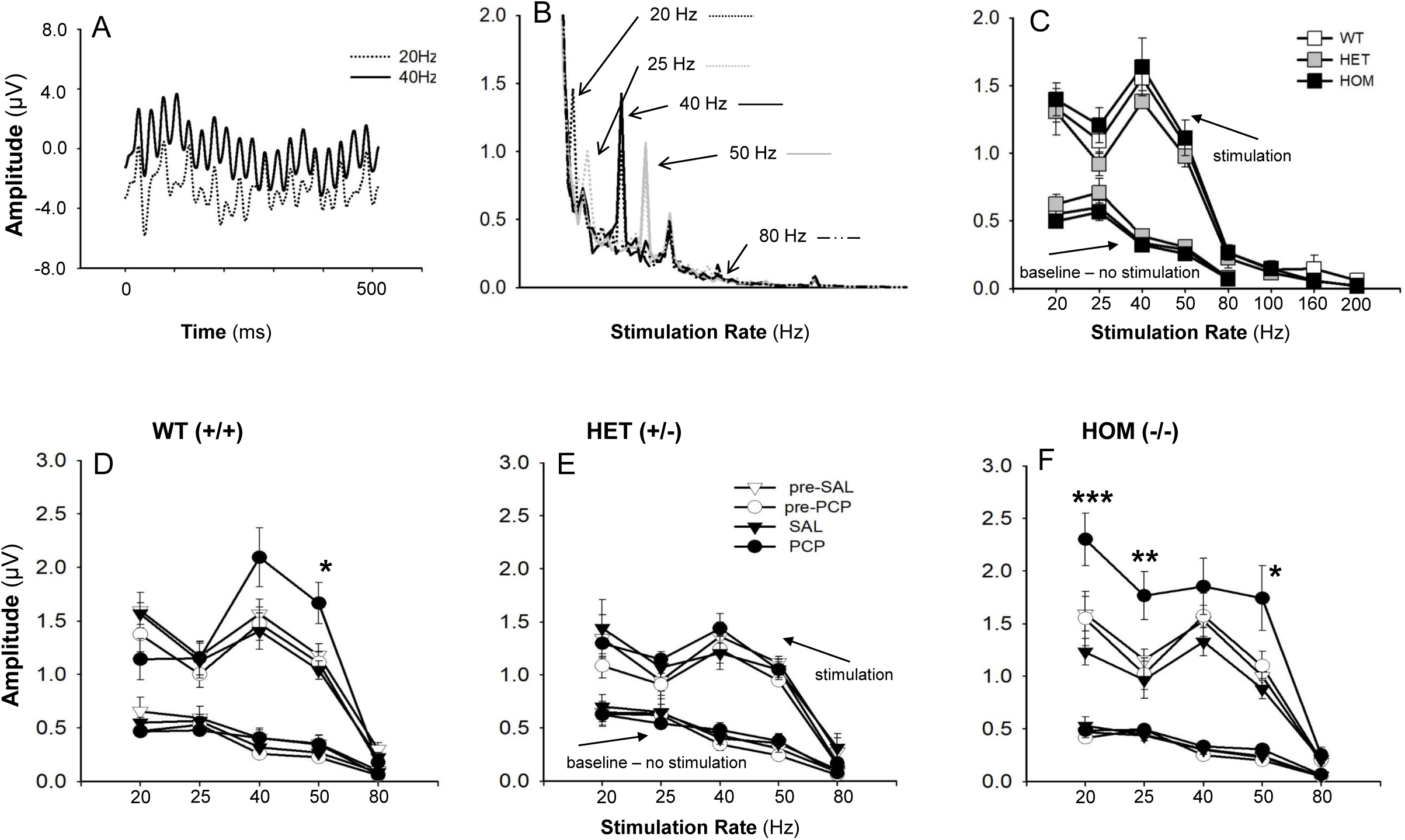
Effect of acute PCP (10 mg/kg) and SAL on auditory steady-state response (ASSR) measured in primary auditory cortex (A1) of SRKO mice. **A:** Responses (amplitude in μV) to 20 and 40 Hz click train stimuli. **B:** Average waveform amplitudes to 20, 25, 40, 50 and 80 Hz responses. **C:** ASSR amplitude by stimulation rate and genotype. Data are the mean ± SEM (n= 14 WT (+/+), n=12 HET (+/-) and n=14 HOM (-/-) from the initial testing day for each animal. **D-F:** Effect of PCP treatment on ASSR amplitude by frequency for WT (**D**), HET (**E**), and HOM (**F**). Data are mean of 7-11 animals per group, across the saline and PCP treatment days. **\*\*\***p = 0.001, **\*\***p ≤ 0.02 and **\***p ≤ 0.05 for saline vs. PCP groups.

#### Pre-stimulation baseline

In the baseline period, there was a significant main effect of frequency (F_4,11_=105.8, p<.0001) F,19=71.3, p<.0001), reflecting a significant linear 1/f reduction in amplitude from 20 to 80 Hz (F_1,22_=312.6, p<.0001) (**Fig 1C**). There were no significant effects of either PCP or genotype on prestimulus (baseline) activity as a function of genotype (**Fig 1C**) or PCP treatment (**Fig. 1D-F**).

#### Stimulation period, initial testing day

In initial testing, the effects of stimulation frequency on power were strongly significant (F_4,132.5_=84.7, p<.0001) with the greatest amplitude occurring at 40 Hz (**Fig. 1C**). The main effect of genotype (F_2,35.9_=1.32, p=.28) on power and genotype X stimulation-frequency interaction (F_8,132.5_=.4, p=.9) were both non-significant.

#### Stimulation period, drug challenge days (**Fig. 1D-F**)

When analyses were conducted across saline and PCP-challenge days, the main effect of stimulation frequency on power was again highly significant (F_4,386.5_=157.9, p<.0001). The treatment day (sal/PCP) X time (pre/post treatment) interaction was also highly significant (F_1,383.2_=25.2, p<.0001) reflecting an increase in ASSR amplitude across all stimulation frequencies and genotypes by PCP.

The main effect of genotype on power was not significant (F_2,19.8_=23.8, p=.083). However, genotype interacted significantly with both stimulation frequency (F_8,386.4_=2.05, p=.04) and drug treatment (F_2,407.6_=7.27, p=.001). The 3-way genotype X time X drug treatment interaction was also significant (F_2,383_=3.70, p=.026), reflecting differential sensitivity to PCP vs. saline across the WT(+/+), HET(+/-) and HOM(-/-) animals.

In order to further parse the significant time X treatment and genotype X time X treatment effect, we calculated difference values for each session reflecting post-vs-pre-treatment values (i.e. within-session drug effects). As expected, the main effect of treatment was again highly significant (F_1,199.7_=42.9, p<.0001) reflecting a larger change during PCP vs. saline treatment. In addition, a significant drug treatment X stimulation frequency effect emerged (F4,182.7=3.18, p=.015). The genotype X drug treatment (F2,199.1=6.38, p=.002) and genotype X drug treatment X stimulation frequency (F8,182.7=3.06, p=.003) effects were both significant.

These final interactions were parsed by conducting separate one-way analyses at each stimulation frequency by genotype and treatment. In WT(+/+) animals, significant PCP effects were observed at 40/50 Hz, but not at other frequencies (**Fig. 1D**). In HET(+/-) mice, no significant PCP effects were observed at any stimulation frequency (**Fig. 1E**). By contrast, in HOM(-/-) mice, significant effects were observed both at lower (20,25 Hz) and higher (40,50 Hz) stimulation frequencies (**Fig. 1F**), reflecting a specific genotype effect primarily on the 20 Hz response (F_2,23_=4.12=.031) with significant post-hoc difference between HOM(-/-) and WT(+/+) animals (LSD p=.009).

### N1 Refractoriness Function

aERP analogous to the human P1/N1 response were obtained in WT(-/-), HET(+/-) and HOM(-/-) SRKO mice in response to tones presented with 1, 3 and 6 sec ISIs. Based upon the ERP we evaluated 3 peaks of interest: P20 was defined as the greatest positivity between 15-35 ms post-stimulus; N40 was defined as the greatest negativity between 30-75 ms post-stimulus; and P80 was defined as the greatest positivity between 70-105 ms (**Fig. 2**). Analogous to ASSR, for each component a 2-way ANOVA with factors of genotype and ISI was conducted on data obtained from the initial testing day for each animal, whereas a 4-way ANOVA with factors of stimulation frequency, genotype, drug treatment (saline vs PCP), and pre-or-post-treatment was conducted across treatment days.

**Figure 2:**
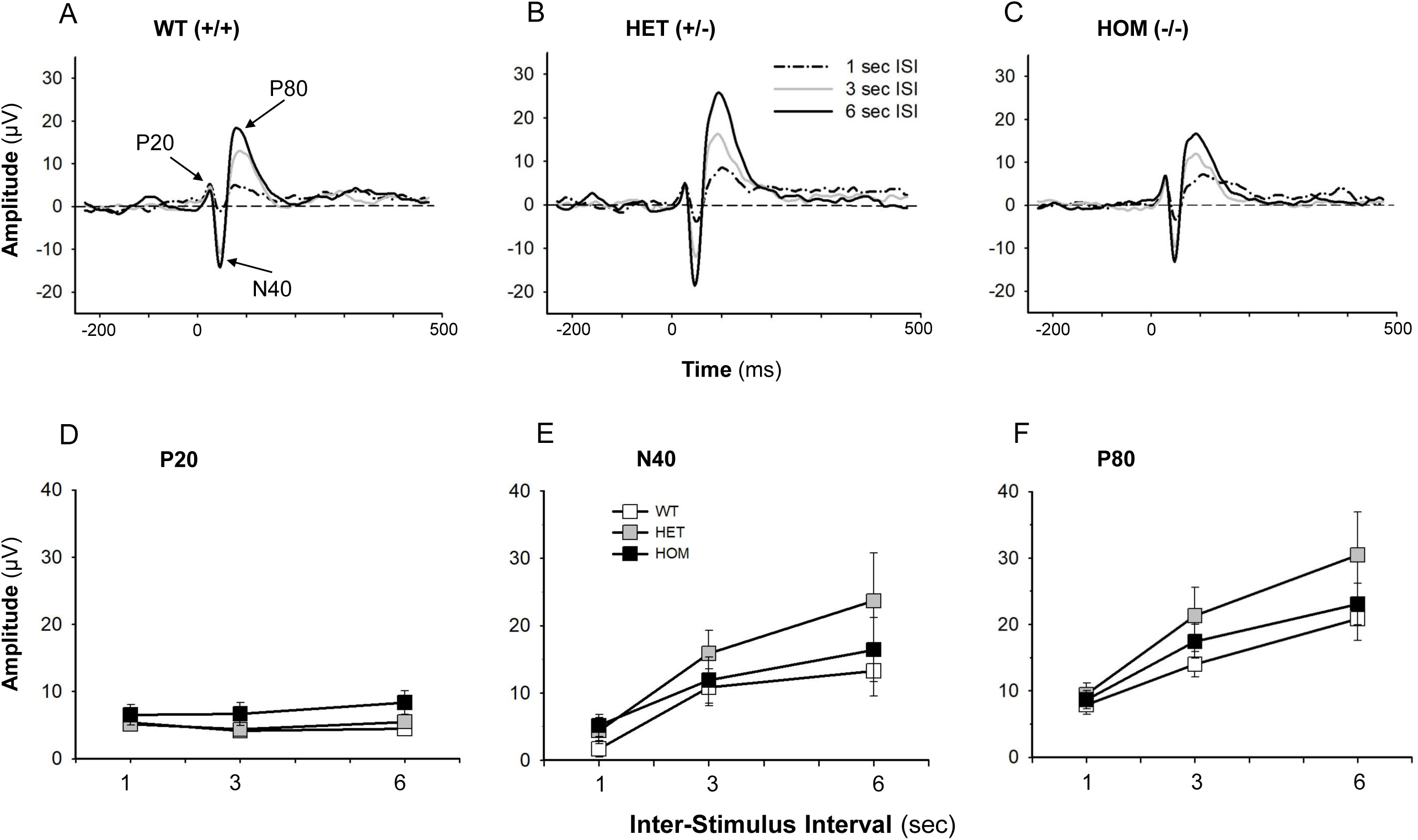
Auditory event-related potentials (aERP) from A1 in response to stimuli presented at 1, 3 and 6 sec in WT(+/+), and HET(+/-) and HOM (-/-) SRKO mice, showing P20, N40 and P80 peaks of interest. **A-C:** aERPs shown as averages of n=12-15 mice. **D-F:** Values are the mean ± SEM (n=11-14).

#### P20

At initial testing, P20 amplitudes were not significantly affected by ISI (F_2,70_=1.31, p=.3) or genotype (F_2,35_=1.64, p=.2). The genotype X ISI interaction was also non-significant (F_4,70_=.8, p=.5) (**Fig. 2A and 2D**).

When analyses were analyzed across treatment day (sal/PCP) and time (pre/post), the ISI effect was again non-significant (F_2,234.3_=2.24, p=.1). The time (F_1,234.3_=.62, p=.4), treatment (F_1,244.5_=.1, p=.7) and treatment X time interaction (F_2,234.3_=.9, p=.4) were all non-significant. The main effect of genotype (F_2,24.1_=2.05, p=.15) and the genotype X time interaction (F_2,234.3_=.1, p=.9) and genotype X treatment interaction (F2,244.0=2.41, p=.09) were also all non-significant as were higher order interactions (**Table 1)**.

**Table 1:**
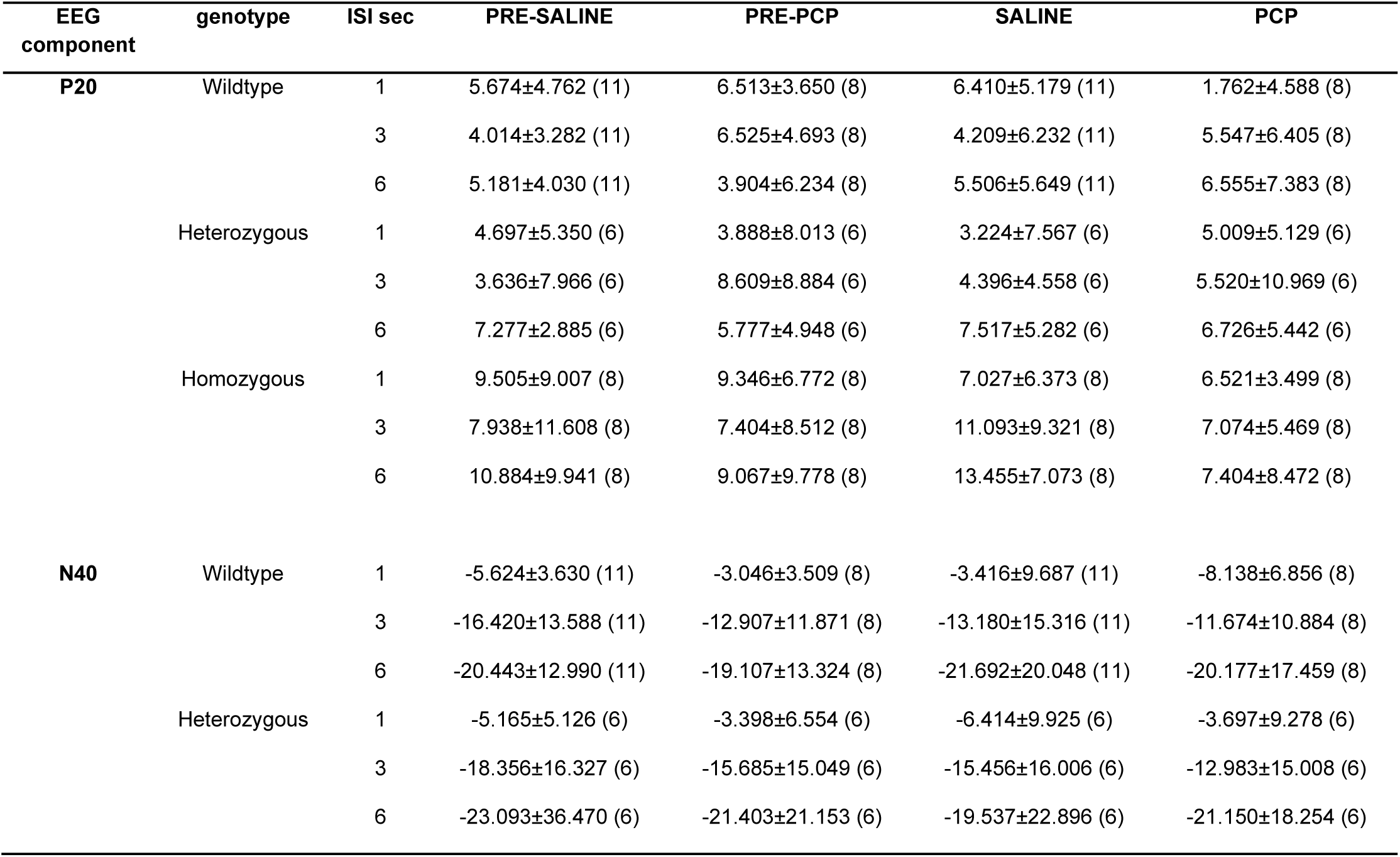
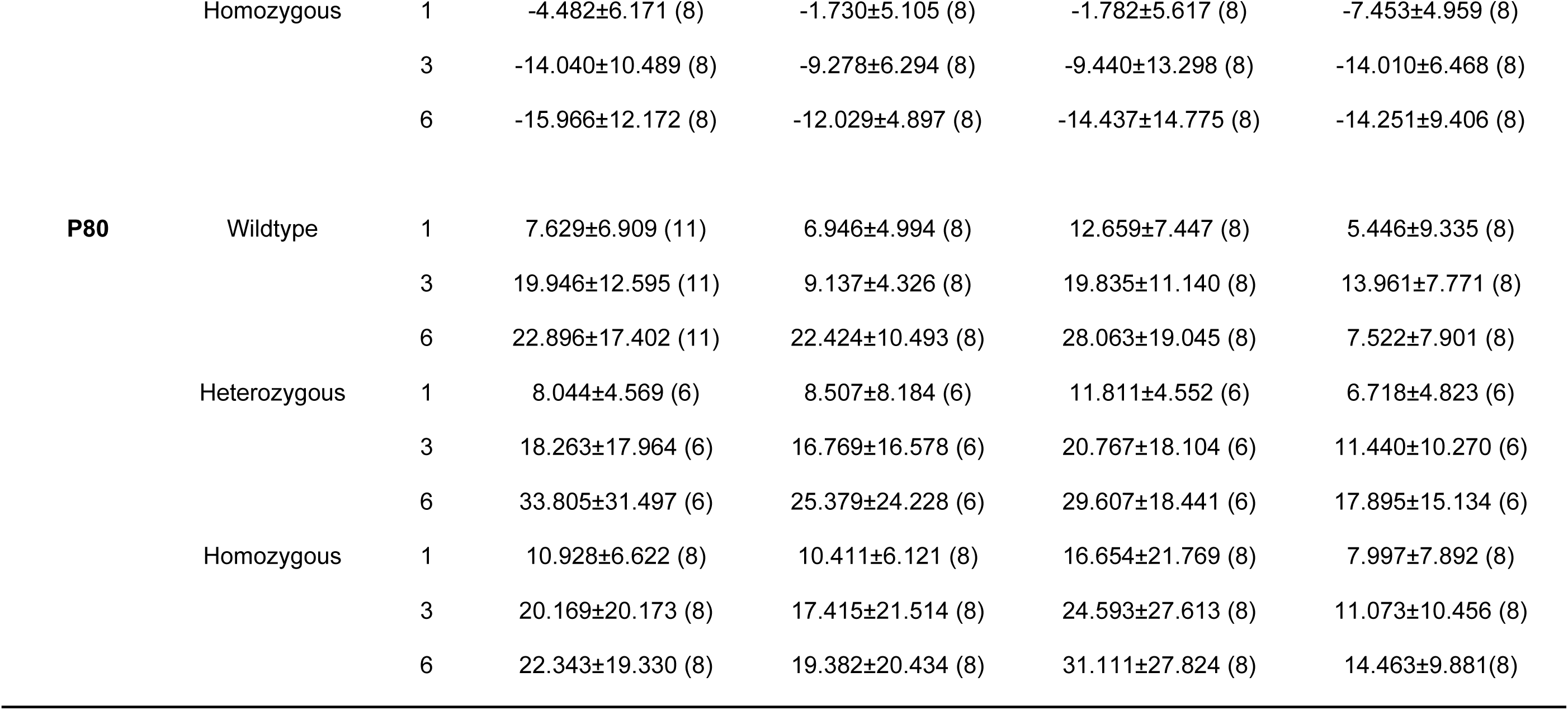
Mean amplitude of indicated aERP components by saline and PCP treatment. Number of animals per observation is shown in parentheses.

#### N40

At initial testing, N40 amplitudes showed a significant ISI effect (F_2,60.8_=31.9, p<.0001) reflecting large N1 potentials with increasing ISI. The effects of genotype (F_2,31.3_=.84, p=.44) and the genotype X ISI interactions (F_4,60.7_=.1.16, p=.3) were both non-significant (**Fig. 2B and 2E**).

When analyses were analyzed across treatment day (sal/PCP) and time (pre/post), the ISI effect was again strongly significant (F_2,233.5_=43.3, p<.0001). The treatment X time interaction was also significant (F_1,233.5_=4.07, p=.045), reflecting a near-significant decrease in N40 amplitude in the PCP vs. saline groups post- (F_1,109.4_=3.89, p=.051), but not pre- (F_1,110.9_=.25), treatment.

The main effect of genotype was non-significant (F_2,23.4_=.33, p=.7), as was the genotype interaction with time (F_2,233.5_=.03, p=.98) and treatment (F2,240.5=.7, p=.5), along with higher order interactions (Table 1).

#### P80

At initial testing, P80 amplitude also showed a highly significant main effect of ISI (F_2,60.9_=45.6, p<.0001), reflecting increased amplitude with increasing ISI (**Fig. 2C and 2F**). The main effect of genotype (F_2,31.3_=1.22, p=.31) and genotype X ISI interaction (F_4,60.8_=1.13, p=.35) were both non-significant.

When analyses were conducted across treatment days, there was a highly significant treatment X time (F_1,234.2_=10.7, p=.001) interaction, reflecting a highly significant reduction in P80 post-vs. pre-PCP (F_1,110.6_=25.7, p<.0001) but not saline (F1,110.0=.64, p=.4). The main effect of genotype was again non-significant (F_2,24.2_=.32, p=.7) as was the genotype interaction with treatment (F_2,234.2_=.24, p=.8) and time (F_2,240.0_=.86, p=.4) along with higher order interactions (**Table 1**).

### Effect of PV specific GRIN1 KO

In order to further evaluate the mechanism underlying increased (rather than decreased) ASSR response following NMDAR antagonist treatment, we evaluated ASSR in an additional group of animals with PV interneuron-specific KO of the GRIN1 subunit. In order to confirm our previously observed NMDAR antagonist effect, we evaluated saline vs. PCP effect in a group of C57/BL6 animals matching the Cre background.

As with studies of the SRKO mice (**Figs. 1D-F**) there was a significant main effect of drug treatment (F_1,47.9_=7.13, p=.01), with PCP increasing ASSR peak power. There was also a trend toward a drug treatment X time interaction (F_1,46.1_=3.77, p=.058). When data were split by pre vs. post treatment no significant difference was observed between drug treatment in the pretreatment period (i.e., pre-saline vs pre-PCP) (F_1,25_=.023, p=.88), whereas a significant difference was observed in the post-treatment period (i.e., post-saline vs post-PCP) (F_1,24.1_=16.1, p=.001), reflecting greater ASSR amplitude in the PCP vs. saline group (**Fig. 3A**).

**Figure 3:**
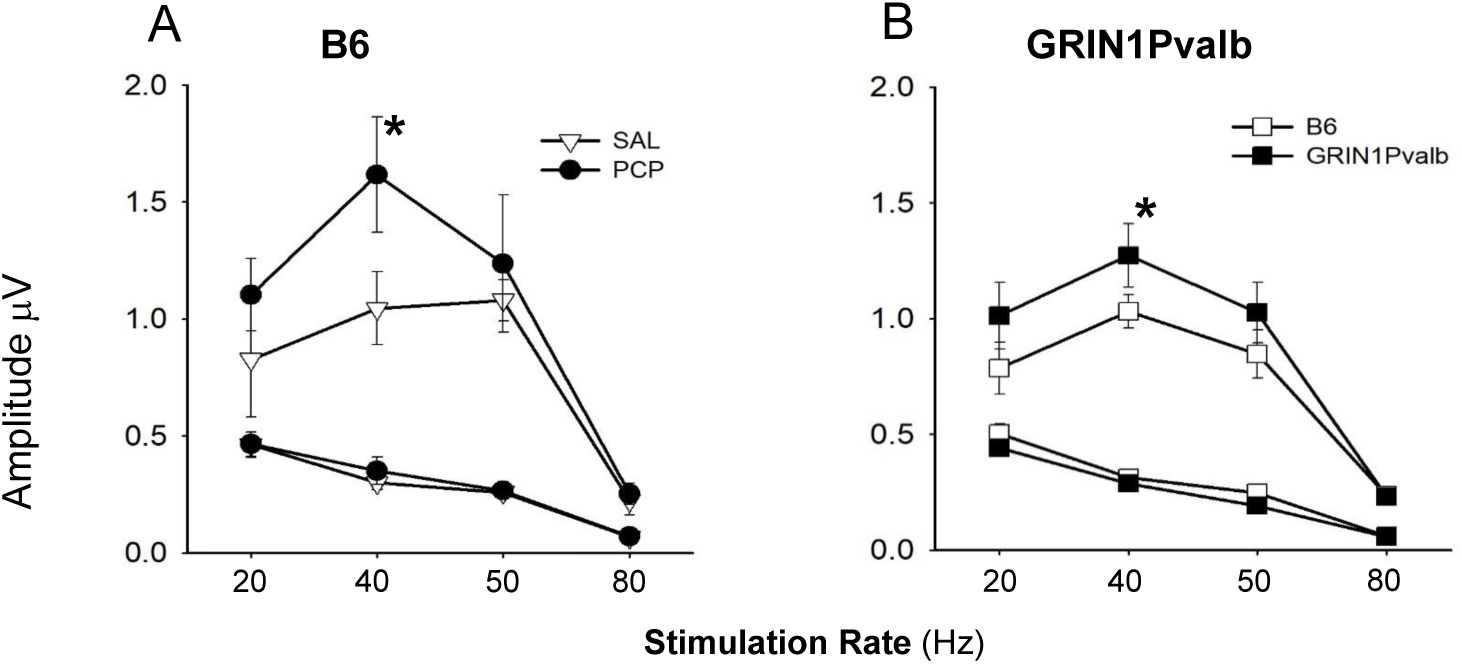
**A:** Comparison of NMDA antagonism by acute PCP (10 mg/kg) versus SAL on ASSR responses (Amplitude in μV) in B6 mice and **B:** in GRIN1Pvalb (NR1 subunit of the NMDAR was knocked-out specifically in PV interneurons) versus B6 mice. Values are the mean ± SEM (n=3-6). **\***p ≤ 0.05 SAL vs. PCP or B6 vs.GRIN1Pvalb mice.

Similar analyses were conducted comparing GRIN1 KO vs. WT animals. Consistent with our acute PCP effects, GRIN1 KO mice showed enhanced ASSR across frequencies (F1,9=5.40, p=.046) as well as at 40 (F1,9=8.26, p=.018) and 50 (F1,9=8.09, p=.019) Hz individually (**Fig. 3B**).

## Discussion

Treatment development research in Sz is hampered at present both by the lack of validated biomarkers that can be used to translate across species, and the lack of animal models that recapitulate key functional features of the disorder. ERP-based neurophysiological biomarkers have become increasingly appreciated given their physiological homology across rodent, monkey and human species. We (Lee et al., 2018) and others (Ehrlichman et al., 2008; Featherstone et al., 2018) have previously investigated the sensitivity of MMN to NMDAR antagonists across species and have documented highly significant homologies in both frequency content and sensitivity to NMDAR antagonist administration.

Here, we evaluated two additional measures – ASSR and auditory N1 refractoriness – that have been shown to be impaired in Sz and may reflect differential involvement of PV- and SOM-related ensembles within rodent auditory cortex. In parallel, we evaluated the degree to which Sz-like phenotypes were reproduced in SRKO mice, a putative model for Sz that is associated specifically with Sz-like PV interneuron downregulation.

Principal findings were that both acute NMDAR antagonist administration (**Figs. 1D, 4A**) and selective NMDAR KO on PV interneurons increased ASSR response (**Fig. 4B**), consistent with the postulated involvement of PV interneurons in ASSR generation. Unperturbed SRKO mice showed ASSR levels that were statistically indistinguishable from WT. However, SRKO significantly modulated the effect of treatment with the NMDAR antagonist PCP, such that HET(+/-) mice showed reduced sensitivity while HOM(-/-) showed significantly enhanced sensitivity (**Fig. 1D-F**).

By contrast to its effects on ASSR, SRKO mice did not significantly affect the rodent N40/P80, which may serve as a rodent analog of the human N1. These findings thus especially support the role of ASSR and N1 as rodent translational biomarkers, and of SRKO mice as an animal model that recapitulates PV-specific aspects of NMDAR dysfunction in Sz.

ASSR has been extensively studied before both in Sz and in rodent models. In Sz, the majority of studies have shown a reduction of ASSR power (e.g., (Hong et al., 2004; Kwon et al., 1999; Light et al., 2006; Reilly et al., 2018; Spencer et al., 2009; Thune et al., 2016), although increases are also reported (Hamm et al., 2012; Kim et al., 2019). Deficits are observed as well across the psychosis spectrum (Parker et al., 2019). Although the reductions have often been attributed to reduced NMDAR dysfunction on PV interneurons themselves, alternative formulations suggest that the high firing rate of PV interneurons is critically dependent upon rapid Ca^2+^ influx through Ca^2+^-permeable AMPA receptors, and that the slower kinetics of Ca^2+^ entry through NMDAR may impede rather than support 40 Hz response (Goldberg et al., 2003; Gonzalez-Burgos and Lewis, 2012).

The inhibitory effects of high-dose NMDAR antagonists on ASSR may thus reflect the additional effects of NMDAR antagonists on current flow within pyramidal excitatory neurons, rather than PV interneurons (Lisman et al., 2008), consistent with our observation that selective NMDAR KO on PV interneurons increases, rather than decreases, ASSR amplitude. These findings are in contrast to findings in mice with KO of NMDAR on more general GABAergic interneurons, where reduction of ASSR is observed (Nakao and Nakazawa, 2014).

Our findings of increased ASSR are consistent as well with other studies of acute NMDAR administration in rodents, which then resolves following chronic treatment (Leishman et al., 2015; Sullivan et al., 2015). Regardless of the precise circuitry of ASSR, however, we demonstrate in this study that SRKO mice show increased sensitivity to the augmenting effects of PCP on the ASSR, supporting that it may serve as an effective model of PV-interneuron dysfunction for translational treatment development.

As opposed to the findings with ASSR, SRKO mice showed unchanged sensitivity to disruptive effects of PCP on the rodent N40/P80 potentials, which may be viewed as a rodent analog of the human N1/P2 potential. Like N1, the rodent N40/P80 shows a prolonged refractoriness function, such that full amplitude is not reached until ISI is prolonged to >6s (Javitt, 2015). In humans, N1 responses map primarily to the theta frequency range, and thus may interrelate with circuit motifs involving non-PV interneurons, especially SOMs (Javitt et al., submitted). Our finding of significant aERP response in mice following NMDAR blockade is consistent with prior studies in monkey (Javitt et al., 2000) and rodent (Bickel et al., 2008; Connolly et al., 2004; Lee et al., 2018; Maxwell et al., 2006). The lack of effect of SRKO on the rodent N1 response may thus suggest that the model recapitulates the PV-related aspects of NMDAR dysfunction in Sz more fully than it captures dysfunction of other neuronal populations.

These findings are also consistent with multi-hit models in which multiple genes contribute in each individual, with each gene only conferring a relatively restricted (1-2%) of the risk (Schizophrenia Working Group of the Psychiatric Genomics, 2014). Here, even complete KO of the SRR gene did not fully recapitulate the NMDAR antagonist-induced phenotype, suggesting that additional genes may also need to be perturbed. At present, relatively little is known about the effects of D-serine on individual interneuron subtypes. Overall, however, these findings are supportive both of use of neurophysiological biomarkers for cross-species comparison, and for KO of genes regulating NMDAR as potential platforms for investigating the effects of different pharmacological agents.

### Limitations

In the present study, we used 3-4 month old mice, which equate to a young adult age range. We consider this age mouse to be most relevant to the pathophysiology of Sz, which typically presents in late adolescence and early adulthood. Nevertheless, pathological consequences of SRKO develop more fully between 3 and 8 months (Coyle and Balu, 2018), suggesting that a more robust phenotype may have been observed in older mice.

Also, in our control experiment we knocked out the GRIN1 subunit of the NMDAR in order to ensure a functional deficit. However, Sz may be specifically associated with disruption of GRIN2A-containing NMDAR (Bitanihirwe et al., 2009; Kinney et al., 2006). Future studies with cell-type specific disruption of GRIN2A and GRIN2B subunits are therefore also required, as well as those targeting other neuronal populations.

Finally, we used only a single acute dose of PCP in the present study, and thus did not confirm the previously reported dose-dependent effects of NMDAR on ASSR (Sivarao et al., 2016). Nevertheless, our dose was sufficient to reduce amplitude of the rodent N1 response, suggesting effective inhibition of NMDAR-mediated neurotransmission.

### Conclusions

Development of NMDAR-based biomarkers and animal models for translational research in Sz remains an important priority area. The present results add to the prior literature that disruption of brain D-serine metabolism affects PV interneuron-related function within cortex as reflected in altered ASSR activity. In addition, we provide additional evidence that PV interneuron-selective NMDAR inhibition increases, rather than decreases, ASSR amplitude in rodents, suggesting that ASSR reductions observed in Sz may reflect dysfunction of NMDAR within other neuronal compartments (e.g. pyramidal neurons, SOM interneurons). We did not observe effects on rodent long-latency aERP components, suggesting that modulation of D-serine metabolism only partially recapitulates the phenotype of Sz. The present results suggest that a combination of aERP measures, including ASSR, auditory N1 and MMN may provide complementary information over the inclusion of any measure in isolation.

## Acknowledgements

We would like to acknowledge the contribution of Dr. Joe Coyle in providing the SRKO mice for these studies.

